# How *Euglena gracilis* swims: flow field reconstruction and analysis

**DOI:** 10.1101/2020.10.12.335679

**Authors:** Nicola Giuliani, Massimiliano Rossi, Giovanni Noselli, Antonio DeSimone

## Abstract

*Euglena gracilis* is a unicellular organism that swims by beating a single anterior flagellum. We study the nonplanar waveforms spanned by the flagellum during a swimming stroke, and the three-dimensional flows that they generate in the surrounding fluid.

Starting from a small set of time-indexed images obtained by optical microscopy on a swimming *Euglena* cell, we construct a numerical interpolation of the stroke. We define an optimal interpolation (which we call synthetic stroke) by minimizing the discrepancy between experimentally measured velocities (of the swimmer) and those computed by solving numerically the equations of motion of the swimmer driven by the trial interpolated stroke. The good match we obtain between experimentally measured and numerically computed trajectories provides a first validation of our synthetic stroke.

We further validate the procedure by studying the flow velocities induced in the surrounding fluid. We compare the experimentally measured flow fields with the corresponding quantities computed by solving numerically the Stokes equations for the fluid flow, in which the forcing is provided by the synthetic stroke, and find good matching.

Finally, we use the synthetic stroke to derive a coarse-grained model of the flow field resolved in terms of a few dominant singularities. The far field is well approximated by a time-varying Stresslet, and we show that the average behavior of *Euglena* during one stroke is that of an off-axis puller. The reconstruction of the flow field closer to the swimmer body requires a more complex system of singularities. A system of two Stokeslets and one Rotlet, that can be loosely associated with the force exerted by the flagellum, the drag of the body, and a torque to guarantee rotational equilibrium, provides a good approximation.

## 1 Introduction

Euglenids are one of the best-known groups of flagellates and are easily found in freshwater [1]. Their biology and cellular structure is well known and over the last decade there has been a growing interest in studying the mechanics underlying their motility. An interesting feature is that Euglenids can exhibit distinct forms of motility (flagellar swimming and amoeboid motion), can switch between them [2], and respond to environmental cues such as confinement and light [3]. This makes them an interesting model system to study sensing and response mechanisms in an elementary, single cell organism.

The amoeboid motion of Euglenids, typically referred to as metaboly, has been studied from an analytical [4], numerical [5] and experimental [2, 6] perspective, and it has inspired the design of of soft robots [7]. The flagellar swimming of *Euglena gracilis*, powered by the non-planar beating of a single anterior flagellum termed “spinning lasso” in the literature, has been studied for a long time but a detailed experimental reconstruction of this complex kinematics has been obtained only recently, and can be found in [8]. It is also known that the organism can modulate the beating of the flagellum to change its trajectory [3], also in response to external light stimuli. In fact, *E. gracilis* is phototactic.

The study of the flow fields induced by unicellular swimmers through their shape changes plays a key role in understanding the interactions of the cell with its living environment [9, 10, 11, 12, 13]. For example, predators may sense their preys through the flows they generate with their motion, or they may induce flows to drive preys towards their feeding organs and capture them (see, e.g.,[14, 15] and the references cited therein). The main features of all these flows can be rationalized by approximating them as the superposition of a few elementary singular solutions [16, 17]. Many different organisms have been analyzed in this way in recent years. Bacteria exhibit mainly rotational flows near the cell body and a pusher flow far away from the cell [16]. As another example, *Chlamydomonas reinhardtii* produces flow fields of different character during its stroke, oscillating between pusher and puller behaviors, while in average it behaves as a puller [18]. The time-dependent features of the flow of sperm cells have been analyzed in [19]. Analyses of this type yield interesting insight in understanding the interactions between different swimmers [20] or between the swimmer and different interfaces [21, 22, 23].

Given the three-dimensional nature of the motion of the flagellum and the lack of obvious symmetries, resolving the flow fields induced by *Euglena* cells would be particularly interesting. The technical challenges involved have prevented this, at least until now. In fact, extending to three dimensions the two-dimensional results on trajectories and flows obtained in recent years is one of the frontiers in the research on the biophysics of micro-swimmers, as testified by the increasing focus on three-dimensional effects in the recent literature [8, 13, 24, 25, 26, 27, 28].

In this paper, we propose to use numerical simulations of the dynamics of swimmer and surrounding fluid to complement the experimental data and overcome some of the difficulties that have limited the study of the swimming behavior of *E. gracilis* so far. Our main results are the following. We provide a reconstruction of the swimming stroke in terms of shapes and of their rate of change and validate it by showing that the application of a hydrodynamic model to the theoretical flagellar waveforms produces swimming trajectories in close agreement with the experimentally observed ones. We further validate our reconstruction of the swimming stroke by computing the three-dimensional flows induced in the surrounding fluid, measuring them with particle tracking velocimetry tecnhiques, and showing good match between the two. Finally, we provide a coarse-grained model of the fluid flows induced by a swimming *Euglena* in terms of a few dominant singularities. A time-varying Stresslet suffices to capture the far field, and the average behavior in one stroke is that of an off-axis puller (i.e., the axes of the puller flow are not aligned with the body axis). Closer to the swimmer body, a system of two Stresslets and one Rotlet is more appropriate to resolve the salient features of the induced flows, loosely associated with the forces exerted by the beating flagellum, the drag of the body, and a torque ensuring rotational equilibrium. In order to capture the fact that the forces exerted by the flagellum at any given time may change character from propulsive to resistive along the flagellum, an even more complex systems of singularities would be required.

A first attempt to reconstruct numerically the swimming behavior of *E. gracilis* has been performed in [8], using a method explored also in [24]. In [8], Resistive Force Theory [29, 30] is used to check the consistency of the experimentally measured trajectories with the motion arising as a consequence of the reconstructed flagellar beat. Quantitative discrepancies were attributed to the limitations of RFT when dealing with hydrodynamic interactions, see [31, 32] for further details. In fact, the complex flagellar kinematics described in [8, 25] requires more accurate models for the nonlocal hydrodynamic forces generated by a flagellum beating in close proximity of the cell body, and in the present work we address the problem using an open source Boundary Element Method [33] that has been validated in a previous publication [32].

The rest of the paper is organized as follows. In Section 2 we confront the following problem: Starting from a set of time-indexed images defining the shapes during a stroke, and obtained by optical microscopy, determine the corresponding history of shape velocities (the rate at which shapes evolve in time). We proceed by numerical interpolation in time and derive an optimized numerical interpolation of the stroke in the neighborhood of each experimental image by minimizing the discrepancy between experimentally measured traslational and rotational velocity of the swimmer and those computed by solving the equations of motion of the swimmer driven by the interpolated shapes. In this way, we obtain pairs (shape, shape velocities) at each time frame, and hence a time-discrete numerical description of the stroke that we call synthetic stroke.

In Section 3, we validate the synthetic stroke by comparing the flow fields it induces, calculated by solving numerically the equations for the fluid flow, with experimental measurements. In order to measure the fluid velocities experimentally, we use the General Defocusing Particle Tracking (GDPT) method [34]. This technique allows for a three dimensional reconstruction of the flow field looking at the displacement of out-of-focus tracer particles with a single-camera view. The defocusing is used to obtain the out-of-plane particles’ position and it is enhanced by using a cylindrical lens to induce a controlled optical aberration in the optical system [35]. We obtain full three dimensional velocity fields and then we use the azimuthal mean introduced in [36] to compare experimental observations with the numerical results obtained with our synthetic stroke.

Finally, in Section 4, we analyze the flow fields induced by the flagellar beating of *E. gracilis* (modeled using our synthetic stroke). We coarse-grain the flow field using Stokes flow singularities. This kind of analysis has been successfully applied to other swimmers such as *C. reinhardtii* [37, 17], sperm cells [19, 38], and bacteria [16, 39]. The procedure extracts the essential characteristics of the flow by approximating it as the superposition of a few singular solutions of the Stokes equations.

We find that, not unlike other organisms [18], the far field is well described by a single time-varying Stresslet. This is confirmed by the analysis of the leading order term of a multipole expansion of the equation describing the flow field, following [40]. A swimming *E. gracilis* cell oscillates between puller and pusher behaviours and its average behavior over one stroke can be idealized as an off-axis puller, i.e., one in which the axes of the puller flow are not aligned with the longitudinal axis of the body.

To coarse-grain the flow field at distances closer to the body of the swimmer we propose a new system of singularities consisting of two Stokeslets and one Rotlet, and we show that this system performs better than other commonly used. The Stokeslets and Rotlet can be loosely associated with the propulsive (and resistive) forces exerted by the flagellum, the drag of the body, and a torque ensuring rotational equilibrium. We also show, however, that the flagellum may exert forces whose orientations and character at any given time may change along its length (e.g., propulsive forces near the proximal end and resistive ones at the distal end). The consequence of this fact is that, to properly resolve the flows induced in close proximity of the swimmer’s body, more complex systems of singularities are needed such as, e.g., multiple Stokeslets along the length of the flagellum.

## 2 Numerical reconstruction of a swimming stroke (synthetic stroke)

This Section aims at constructing a numerical representation of a swimming stroke starting from the knowledge of a finite set of time-indexed images from optical microscopy. The idea is to interpolate in time the experimental images of the swimmer and of its flagellum and to choose an optimal interpolation capable of reproducing accurately the motion of the swimmer and the induced flows. We call this optimal interpolation synthetic stroke. This is obtained by minimizing the discrepancy between experimentally measured translational and rotational velocities of the swimmer and those computed by solving the equations of motion of the swimmer driven by the interpolated stroke.

The procedure we adopt is based on two key steps, namely, (i) the mathematical definition and numerical solution of the swimming problem, and (ii) an optimization procedure to extrapolate flagellar shapes to pairs of flagellar shapes and shape velocities. The procedure leads to the definition of a numerical time-discrete synthetic stroke starting from a limited set of experimental observations.

### 2.1 Mathematical formulation of the swimming problem

We follow [41, 42] to represent a model swimmer as a time-dependent bounded open set *B*_*t*_ ∈ ℝ^3^. We describe the motion using the map 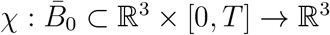 which carries a material point **X** of the swimmer into its current position **x** at time *t*, namely

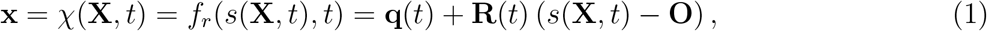

where bold symbols denote points, vectors, and tensors. According to equation (1), *χ* results from the composition of prescribed shape changes *s*(**X***, t*) and a rigid motion *f*_*r*_ of translation **q**(*t*) and rotation **R**(*t*). We set *B*_*t*_ = *χ*(*B*_0_*, t*), see Fig. 1. As for the velocity of any material point on Γ = *∂B*_*t*_, from (1) we obtain

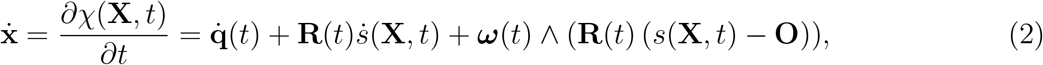

where we have introduced the angular velocity ***ω***(*t*). Assuming that shape changes are prescribed through a periodic function of time, the unknowns of the swimming problem are 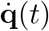 (*t*) and ***ω***(*t*), which characterize the translational and rotational motion of the swimmer (referred to as the rigid velocities in what follows).

**Figure 1:**
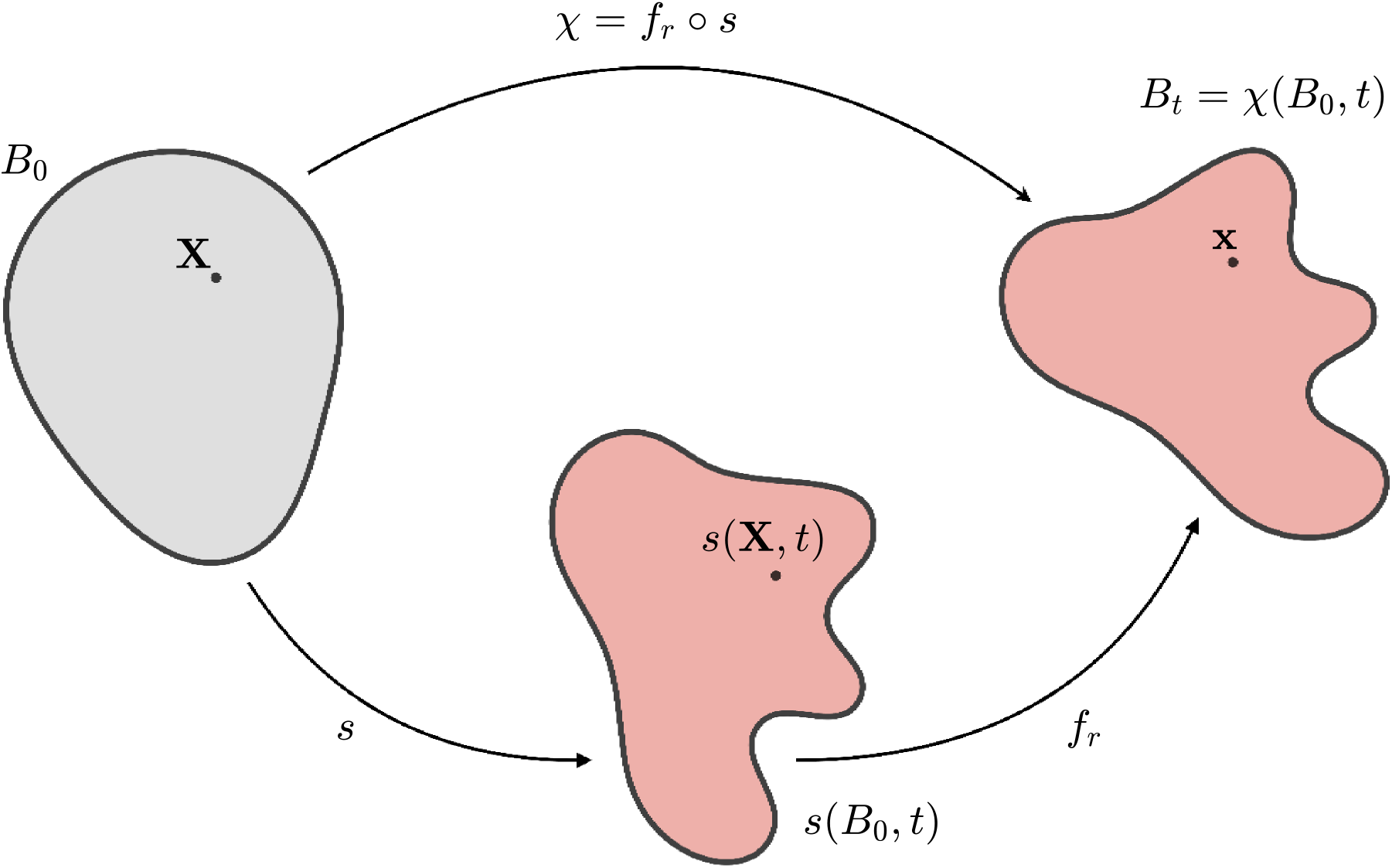
A sketch of the swimmer motion *χ*(**X***, t*) to highlight its decomposition into prescribed shape changes *s*(**X***, t*) and a rigid motion *f*_*r*_.

In view of the characteristic length and time scales relevant to micro-swimmers, inertial effects are negligible such that the balance of linear and angular momentum read

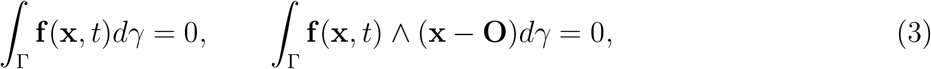

where **f**(*x, t*) is the viscous traction acting on the boundary of the swimmer. This is given by the action of the Cauchy stress tensor ***σ*** [43], namely

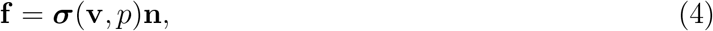

where **n** is the outer unit normal to the boundary Γ, and (**v***, p*) represent the velocity and pressure fields in the fluid. For an incompressible Newtonian fluid the Cauchy stress tensor is given by

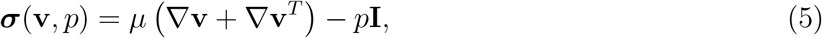

so that, in the low-Reynolds number approximation of micro-hydrodynamics, fluid flow is governed by Stokes’ equations [44]. Neglecting gravity (here we consider a neutrally buoyant swimmer) and assuming that the swimmer moves in free space, these read

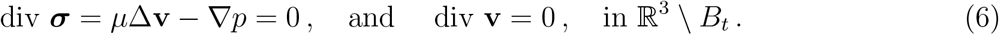

These are complemented by no-slip boundary conditions at the swimmer boundary, where **v** must match the velocity of the swimmer given by (2), and decay conditions at infinity. Notice that the linearity of Stokes’ equations leads to a linear dependence of the tractions **f**(**x***, t*) appearing in (3) on the shape velocities *ṡ*(**X***, t*) (data) and on the rigid velocities 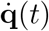 and ***ω***(*t*) (unknowns) appearing in (2). Solving (3) for the latter, we obtain the instantaneous translational and rotational velocities of the swimmer resulting from shapes changing at rate *ṡ*.

We solve numerically the Stokes system by exploiting a Boundary Element Method, and among the different possible implementations we follow [45, 33]. Figure 2 shows the geometric BEM discretization of *Euglena* consisting of 1032 different cells. Further information about the numerical scheme are available in Section I of the supplementary material.

**Figure 2:**
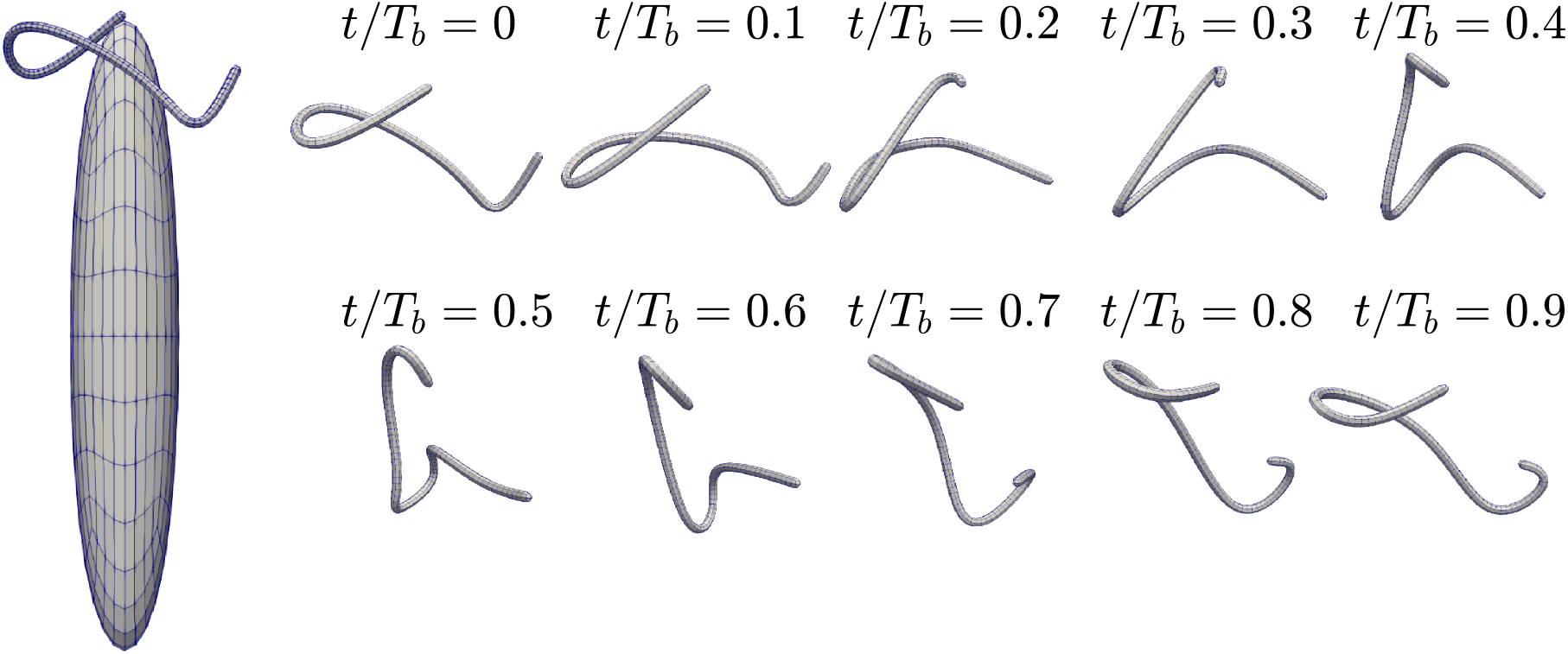
Geometric reconstruction of *E. gracilis*: 384 cells constitute the cell body and 648 the flagellum. We report on the right the 10 flagellar shapes corresponding to the 10 times *t* used to discretize the beating period *T*_*b*_ in [8].

### 2.2 Optimal numerical interpolation based on experimental rigid velocities

The numerical procedure exploiting the BEM defines a unique map between the function giving the history of the shapes of the swimmer during one stroke and its rigid velocities (translational and rotational) during the stroke. In principle, we could write

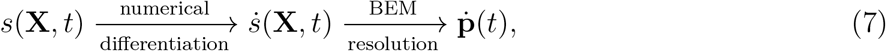

to obtain the rigid velocities 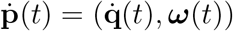 from prescribed *s*(**X***, t*).

In practice, information on the swimmer shapes is only available through finitely many snapshots coming from optical microscopy. For example, the experimental observations of [8] provide ten flagellar shapes during one stroke. However, as shown in Section 2.1, the rigid velocities of the swimmer, **ṗ** (*t*), can be computed by solving the equations of motion (3) only if the rate at which flagellar shapes change, *ṡ*(**X***, t*), is known. The experimental observations of [8] do not provide this information, and we discuss now how we overcome this problem, by finding a suitable interpolation in time of the ten experimentally measured shapes.

Clearly, there are infinitely many interpolating paths. We reduce the degeneracy by exploiting the knowledge of the experimental values for the rigid velocities at the ten instants at which flagellar shapes are known (Figures 5F 5G of [8]), and by defining an optimal interpolation that minimizes the discrepancy between experimentally measured translational and rotational velocities of the swimmer and those computed by solving the equations of motion (3) of the swimmer driven by the interpolated stroke. In this way, we obtain the pairs (shape, shape velocities) at each time frame, and hence a time-discrete synthetic stroke. The procedure is described in more detail below.

We consider the ten flagellar shapes 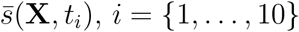, reported in [8] and we interpolate them in time using spline interpolation. To this end, we introduce spline interpolants *g*_*j*_(**X***, t*) with degree *j* = {1*, …, N*_*g*_}. We fix the maximum degree *N*_*g*_ = 5 to obtain a good interpolation while limiting oscillating phenomena related to high order polynomials [46]. We reconstruct shapes as linear combinations of the *N*_*g*_ basis splines in a neighborhood of the ten experimental frames, namely

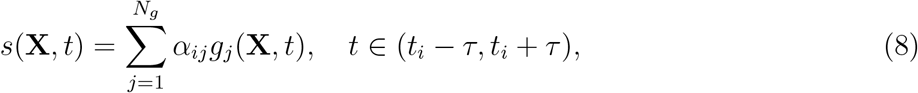

where *α*_*ij*_ are coefficients to be determined by best fit of the experimental data, and *τ ≪ t*_*i*+1_ − *t*_*i*_ specifies the neighborhood of the experimental frame. To ensure that (8) interpolates the experimental flagellar shapes we require that 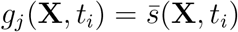 for all *j*, which is possible since 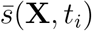 is given in [8] as a spline, and that

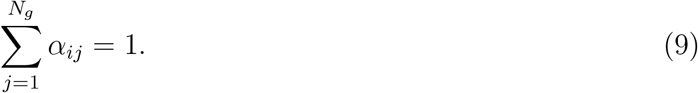

We notice that

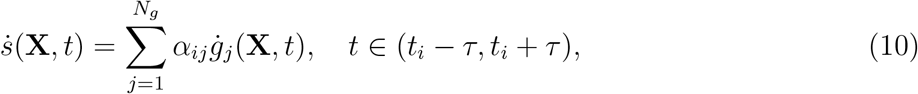

and that we can associate some rigid linear and angular velocities to the basis shape velocities *ġ*_*j*_(**X***, t*),

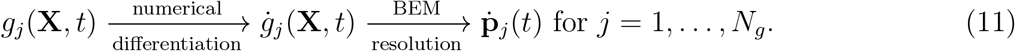

By the linearity of the Stokes system it follows from (10) that

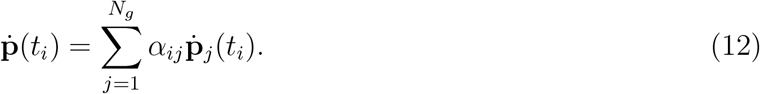

To determine the coefficients *α*_*ij*_ of the optimal interpolation (8) we require that

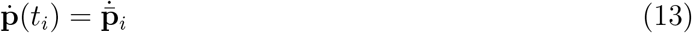

where 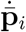 is the value measured experimentally in [8] at the specific frame *t*_*i*_. Notice that the set of experimental data comprises 6 × 10 rigid velocities and we use 5 different coefficients for 10 different time-frames. We exploit a constrained least square minimization algorithm [47] to solve this over-determined system and find values of *α*_*ij*_ fulfilling (9).

The results of the fitting procedure are shown in Figure 3: (a) reports the linear velocities, whereas the angular velocities are shown in (b). Red, green, and blue curves represent their components along the unit vectors of the body reference *i, j, k*. We plot with stars the results of the optimal spline interpolation and we compare them with the reconstruction of [8], represented with triangles. Remarkable agreement is found between our current methodology and the reference results. As for the components *ω*_*x*_ and *ω*_*y*_ of the angular velocity, these are recovered with slightly less accuracy. This minor difference is due to the fact that those two components are the ones with the lowest magnitude and consequently are less relevant in the least square resolution of (13) with constraint (9).

**Figure 3:**
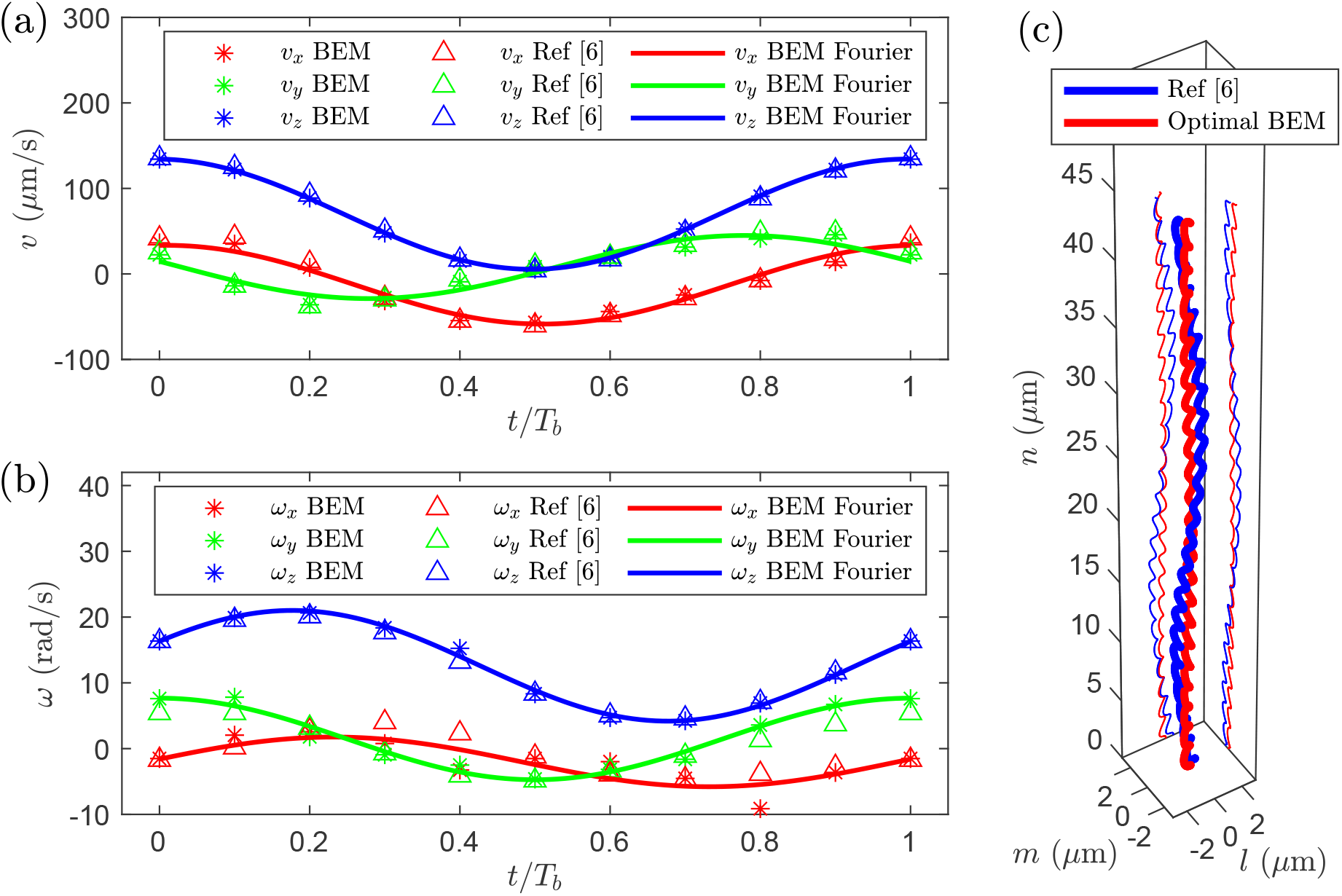
Results of the fitting procedure: (a) and (b) represent linear and angular velocities, respectively. Stars and triangles are used to represent the BEM optimal results and the reference experimental reconstruction of [8]. Continuous lines are the Fourier continuation of the BEM optimal results we use to compare the trajectories in (c). Red, green, and blue refer to the component along *i, j, k*, respectively. (c) comparison between the reference trajectory of [8] (in blue) and the trajectory from the optimal BEM solution (in red) in a reference frame whose vertical axis is aligned with the average direction of motion.

In order to define the trajectory associated with the synthetic stroke, we construct a continuous approximation of 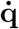 and ***ω*** by using a Fourier expansion of the BEM data. We remark that this is consistent with the procedure used in [8] for the reconstruction of the experimental data. The continuous approximations are shown as continuous lines in Figures 3(a)-(b). We use a standard numerical integration to get the trajectory associated with our synthetic stroke (giving the optimal numerical time-interpolation of the observed swimmer shapes) and we compare it with the experimental observations of the real swimmer in Figure 3(c). Good agreement between our numerical reconstruction and the experimental observations is found also in terms of integrated trajectories.

Once the pairs (shape, shape velocities) defining a time-discrete synthetic stroke are known, one can use the BEM-based algorithm not only to compute translational and rotational velocity of the swimmer, but also to evaluate the velocity field at any point of the fluid domain ℝ^3^ \ *B*_*t*_. We remark that the presence of physical walls, needed in the experimental setting, does not seem to play a very significant role in the current analysis. In fact, *Euglena* swims with its longer body axis parallel to the walls (slide and coverslip) and in the transversal direction the diameter of the body (9*μm*) is one tenth of the channel width (80*μm*). At these distances, the flow velocity computed form the free–space solution has decayed to 10% of the average swimming velocity (the rigid translational velocity). To double-check that the presence of the walls does not alter significantly the conclusions of our analysis, we have computed the rigid velocities associated with our synthetic stroke in the presence of two no–slip walls, located at the same distance as in experimental setting, and we have found that the velocities variations over the stroke are, depending on the velocity component, between 1.7% and 16.5%. We conclude that, in agreement with [8], walls do not play a major role in the observed swimming behavior of *Euglena*.

In what follows, we use the flow velocity fields computed numerically from our synthetic stroke for further validation and analysis. In particular, in Section 3 we use the comparison of fluid flow velocities measured experimentally with those calculated from the synthetic stroke to confirm the reliability of the synthetic stroke. In Section 4 we use the flow fields reconstructed numerically from the synthetic stroke to extract the main qualitative features of *Euglena*’s swimming stroke.

## 3 Experimental validation of the synthetic stroke

As discussed earlier, BEM computations allow for the determination of the three-dimensional flow field induced by a swimming *Euglena* at given instants in the stroke with a degree of detail which is difficult to achieve experimentally. Nevertheless, an experimental validation of the velocity fields obtained numerically is required to validate the results of a computational model. Conventional 2D Particle Tracking Velocimetry (PTV) has been used in the past to measure the flow field around micro-swimmers [48, 14, 49]. Due to its inherent limitations, this technique allows to measure only the components of the velocity parallel to the focal plane, with no information about the azimuthal velocity component [36]. To overcome these limitations, we rely here on the General Defocusing Particle Tracking (GDPT) method [34] to measure the full 3D flow field around a swimming *Euglena*. To the best of our knowledge, this is the first time that 3D PTV measurements around a swimming micro-organism of this kind have been performed. To compare with the numerical results, we follow the approach proposed in [36] and consider time- and azimuthally-averaged velocity fields, but this time accounting for all the velocity components, *i.e.* the radial, the vertical, and the azimuthal component. From now on we indicate with *x, y, z* the coordinates in the body reference frame identified by three unit vectors *i, j, k*, and with *x*′*, y*′*, z*′ the coordinates in the laboratory frame.

### 3.1 Velocity measurements using General Defocusing Particle Tracking (GDPT)

GDPT is a single-camera method that allows to measure the position in space of monodisperse tracer particles observed with an optics with small depth of field [50]. The corresponding particle images recorded with such a systems show distinctive defocusing patterns that depend on the particle position along the optical axis. To enhance the shape deformation and break the symmetry of the defocusing function along the depth direction, we used a cylindrical lens in front of the camera sensor to introduce a mild astigmatic aberration in the optical system [51, 35]. GDPT is based on a look-up table approach to classify the particle-image defocusing patterns in relation to the particle depth position. The normalized cross-correlation is used to rate the similarity between target particle images and the reference images in the look-up table, and hence determine their respective depth position [52]. An exemplary micrograph with defocused particle images observed with the astigmatic optics and the respective 3D particle position reconstruction performed by GDPT is reported in Figures 4(a)-(b). Clearly, particles in the image region occupied by the cell body could not be processed. The tracer particles are polystyrene beads with diameter of 1 μm (Life Technologies, catalog number F8821) at low concentration (volume fraction of 0.06 %). A complete description of the experimental setup can be found in the Supplemental Material.

**Figure 4:**
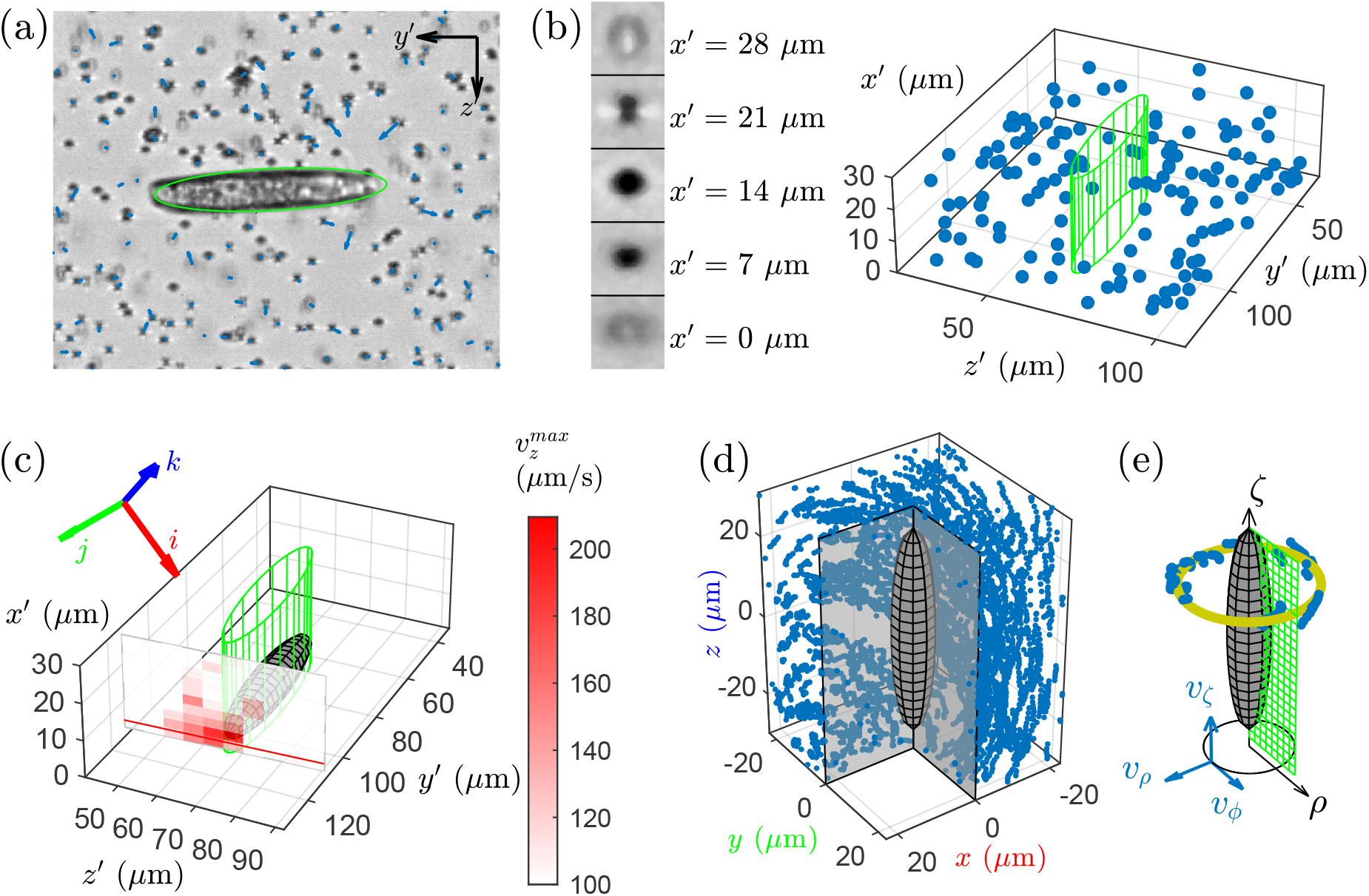
(a) micrograph of a swimming *Euglena* surrounded by tracer particles obtained with 64× magnification and astigmatic optics. (b) reconstruction of 3D particle positions using the GDPT method. The depth position of tracer particles is determined from the different shape of the defocused-astigmatic particle images. (c) determination of the depth position of the swimmer by looking at the maximum axial velocity of the fluid in the swimmer wake. (d)-(e) determination of the time- and azimuthally-averaged velocity components. All particle positions and velocities are reported in the coordinate frame of the swimmer (*ijk*) and converted in cylindrical coordinates. The time- and azimuthally-averaged velocity components *v*_*ρ*_, *v*_*ζ*_, and *v*_*ϕ*_ are calculated as a function of *ρ* and *ζ*.

**Figure 5:**
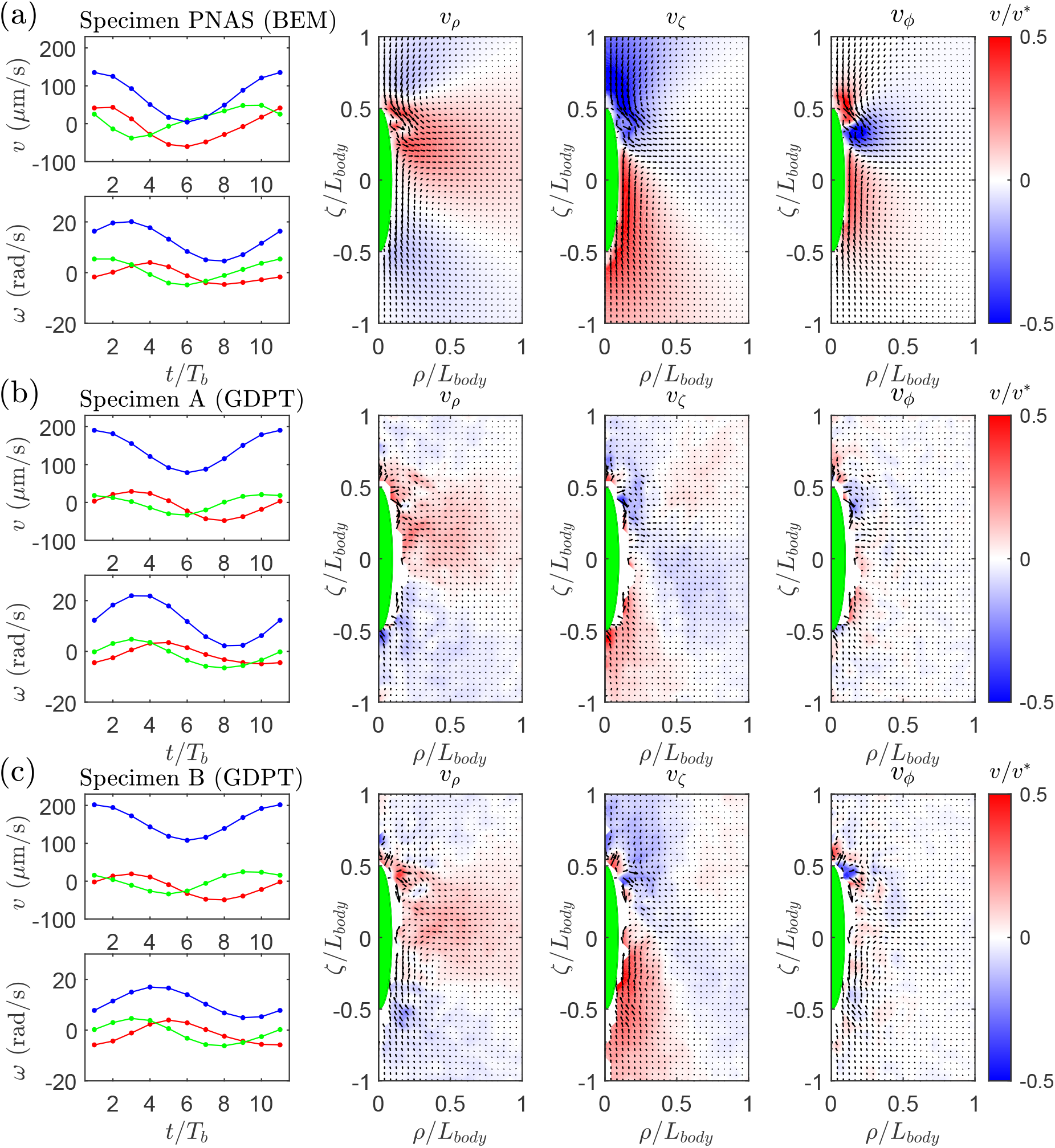
Fluid flow comparisons in which each row shows results from a different specimen. The first column reports the body linear and angular velocities (red, green and blue represent *x, y, z* components in body frame). The remaining three columns depict the different axisymmetrically projected velocities *v*_*ρ*_, *v*_*ζ*_, *v*_*φ*_. The colormap is normalized using the mean velocity of the swimmer during the stroke *v*^*^.

With the current setup we obtained a measurement volume of 168 × 147 × 28 μm^3^, with an estimated uncertainty in the determination of the particle position of 0.02 μm (0.15 pixels) for the in-plane coordinates and of 0.2 μm for the out-of-plane coordinate. As for the uncertainty in the velocity, this is affected by several factors: the particle positioning error, the time interval between two images (1 ms), and the Brownian motion of the particles. We estimated this uncertainty looking at particles in regions of stagnant fluid, obtaining values of 30 μm/s for the in-plane velocity components and of 120 μm/s for the out-of-plane component. These are of the order of magnitude of the velocities to be measured. For this reason, it was not possible to achieve time-resolved velocity fields and we had to consider smoothed time- and space-averaged data.

For each experimental recording, the 3D trajectory and orientation of the cell was obtained using the procedure described in [8], except for the depth position. In fact, this was determined by looking at the maximum axial velocity component 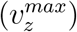 of the flow field in a plane perpendicular to the cell body axis and just behind it, see Figure 4(c). Under the approximation that the cell was swimming parallel to the image plane, the depth position could be identified by the point of maximum average velocity. With the knowledge of cell position and orientation, the velocity vectors measured by GDPT were reported in the reference frame of the cell, Figure 4(d), so that the cylindrical components of the flow velocity field around a swimming cell could be computed as defined in Figure 4(e). In the figure, *v*_*ζ*_ denotes the vertical component, *v*_*ρ*_ the radial component, and *v*_*ϕ*_ the azimuthal component. Finally, the time- and azimuthally-averaged velocity components were reported on a regular grid with pitch 1/48 of the swimmer length (pitch size of approximately 1 μm). At each point of the grid, velocity components were computed as based, on average, on 40 velocity samples, leading to a final estimated uncertainty of about 5 μm/s in the radial and vertical direction and of 18 μm/s in the azimuthal direction.

### 3.2 Comparison between experimental and numerical velocity flow fields

We report in Figure 5(a) a comparison between the numerical reconstruction and the experimental results for the average flow field around the swimmer body. We represent two different experimental specimens to better identify the characteristic structures of the flow. It should be noted that a thorough analysis of the variance of flagellar beating geometries of *E. gracilis* specimens was performed in [8]. Such analysis showed that, beside some variability in the kinematic parameters, there is no substantial differences in the beating patterns of different specimens.

The first column in the figure shows the linear and angular velocities of the cell body within a stroke. Clearly, the results relative to the specimen from [8], used as reference for the numerical reconstruction of Section 2, and those from the experimental observation of other two samples share common distinctive features. For instance, the different components of linear and angular velocity appear in phase, a fact that allows for the comparison of velocity fields from different specimens. This is reported in the other columns of the figure in terms of the velocity components relevant to the axisymmetric projection: *v*_*ρ*_, *v*_*ζ*_, *v*_*ϕ*_. In the three columns, we identify structures that appear to be characteristic of the average flow field around a swimming *Euglena*. Specifically, all the scalar fields can be subdivided in three different regions of alternating positive and negative sign. Furthermore, the fields in the second and third columns (*i.e.*, those for *v*_*ρ*_ and *v*_*ζ*_) clearly exhibit the behavior of a puller swimmer: the fluid is pulled by the swimmer at the top (negative *v*_*ζ*_) and at the bottom (positive *v*_*ζ*_), while it is pushed radially at the center (positive *v*_*ρ*_). We note, in passing, that some differences emerge with respect to a typical puller, which is characterized by zero longitudinal velocity at *ζ/L*_*body*_ = 0. The fact that, in the current analysis, *v*_*ζ*_ is different from zero (mainly negative) at these locations suggests that the pulling forces are at an angle with respect to the body axis of the swimmer. This feature, which is due the intrinsic lack of symmetry in the flagellar beat of *Euglena*, was already observed in [49]. We will return to it in the next section.

The component *v*_*ϕ*_ of the velocity field (fourth column in Fig. 4) depends strongly on the rotation of the swimmer about its major axis. This can be described as a counter-clock wise rotation around the body axis oriented from the posterior to the anterior end. We identify two positive regions at the top and at the bottom, and this observations agrees with the direction of rotation of the swimmer. Moreover, the asymmetric beating of the flagellum induces a rotational flow localized at the flagellum, and rotations of the body of opposite sign, which in turn induce a counter-rotating flow around the body. The negative tangential velocity in the central part of the body is consistent with the existence of this counter-rotating flow.

While the measured components of *v*_*ρ*_ and *v*_*ζ*_ compare well with the corresponding computed fields, a comparison for the component *v*_*ϕ*_ is more delicate. To understand why, we recall that Stokes flow can be coarse-grained by using fundamental solutions of the Stokes system; the basic Stokes singularity associated with rotations is the so-called Rotlet, which induces a quadratic flow decay with the distance from the location of the singularity. Since the beating of the flagellum leads to the presence of two counter-rotating flows, the velocity component *v*_*ϕ*_ exhibits a cubic decay (we will investigate this aspect in more detail in Section 4.1). This rapid decay leads to a very low intensity of the signal to be measured. This fact, combined with the lower resolution of GDPT for the out-of-plane velocity component with respect to in-plane ones, implies a lower signal to noise ratio in the field *v*_*ϕ*_ with respect to the fields *v*_*ρ*_ and *v*_*ζ*_. In spite of this, we are still able to identify the three different regions highlighted by the computational analysis, confirming that the flow structures identified by the BEM are characteristic of the real flow field around a swimming *Euglena*.

We conclude that the synthetic stroke defined in Section 2 approximates well not only the rigid body kinematics (linear and angular velocities, and trajectories of the cell body) but also the essential characteristics of the flow fields induced in the surrounding fluid.

## 4 Flow analysis using singularity approximations

We move now to the analysis of the flow field induced by a swimming *Euglena*. In particular, we first introduce possible approximations for the velocity field of the synthetic stroke as based on a combination of Stokes flow singularities and then discuss the swimming behavior of *E. gracilis*.

### 4.1 Coarse graining the flow field using singularities

Typically, micro-swimmers exert complex systems of forces on the surrounding fluid. From a physical perspective, these are described by the traction **f** at the swimmer boundary – recall the balance equations (3) – and can be computed using the BEM mentioned in Section 2. While this is the exact, infinite-dimensional parametrization of the forces exerted by a swimmer, it is often of interest to find a statically equivalent, finite-dimensional system of singularities (i.e., a system of finitely many concentrated forces and torques) capable of reproducing the main features of the surrounding fluid flow with sensible accuracy. In this way, one can arrive at a coarse-grained representation of the swimmer, and of the flow it induces, which is of great help in formulating a compact, conceptual picture of its interactions with the environment. This is the purpose of the present section, where we introduce and compare different singularity models for a swimming *E. gracilis*.

The flow field around micro-swimmers has often been approximated by a combination of fundamental solutions of Stokes flow. Applications of this approach range from swimming sperm cells, represented in [19] by means of force singularities (called Stokeslet), to swimming bacteria, whose flow field has been coarse-grained through a combination of two Stokeslet (Stokeslet doublet) and two torque singularities (Rotlet doublet), see [16, 53]. As an example of particular interest for the present study, the reconstruction of the fluid flow induced by *C. reinhardtii* has been successfully achieved in [36] by means of a system of three Stokeslets.

In view of the balance equations (3), we require the system of forces, **F**_*i*_, and of torques, **T**_*i*_, representing the actions of a swimmer on the surrounding fluid to satisfy

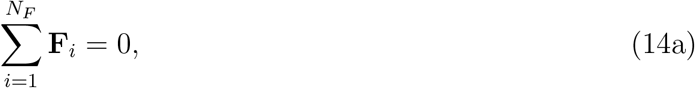

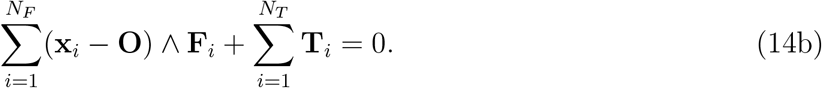

In order to find a reliable coarse-grained model for the flow field induced by *E. gracilis* in terms of a limited number of singularities, we consider the values of the fluid velocity as computed using the BEM on spherical shells surrounding the swimmer with internal and external radii *R*_*min*_ and *R*_*max*_, respectively. We use three different ranges to test the singularity approximations: nearby points between *R*_*min*_ = 1.5*L*_*body*_ and *R*_*max*_ = 5*L*_*body*_, intermediate points between *R*_*min*_ = 5*L*_*body*_ and *R*_*max*_ = 15*L*_*body*_, and faraway points between *R*_*min*_ = 15*L*_*body*_ and *R*_*max*_ = 25*L*_*body*_. We denote here by *L*_*body*_ the characteristic length of the swimmer (~ 50 μm), and consider a regular grid in spherical coordinates using 11 points along the radius, the polar and the azimuthal angle, for a total number of 1331 points for each spherical shell.

Given the constraints on forces and torques as expressed by (14), the simplest system of singularities consists of two directly opposing forces (Stokeslet doublet, which originates the Stresslet singularity when the distance between the points of application of the forces tends to zero). We provide in Table 1(a) a sketch of this configuration. Comparing with the situation arising in the case of a swimming bacterium, we recall that this configuration well reproduces the far field velocity while a satisfactory agreement also for the near field velocity is achieved only by including two opposing torques (Rotlet doublet) to account for the rotational flows induced by the rotational motion of the flagellum and by the counter-rotations of the body, see [53] for more details. The additional Rotlet doublet does not significantly perturb the far field because the flow field induced by a Stokeslet doublet decays as *r*^−2^ while the one induced by a Rotlet doublet decays as *r*^−3^ (here *r* is the distance from the location of the singularity).

**Table 1:**
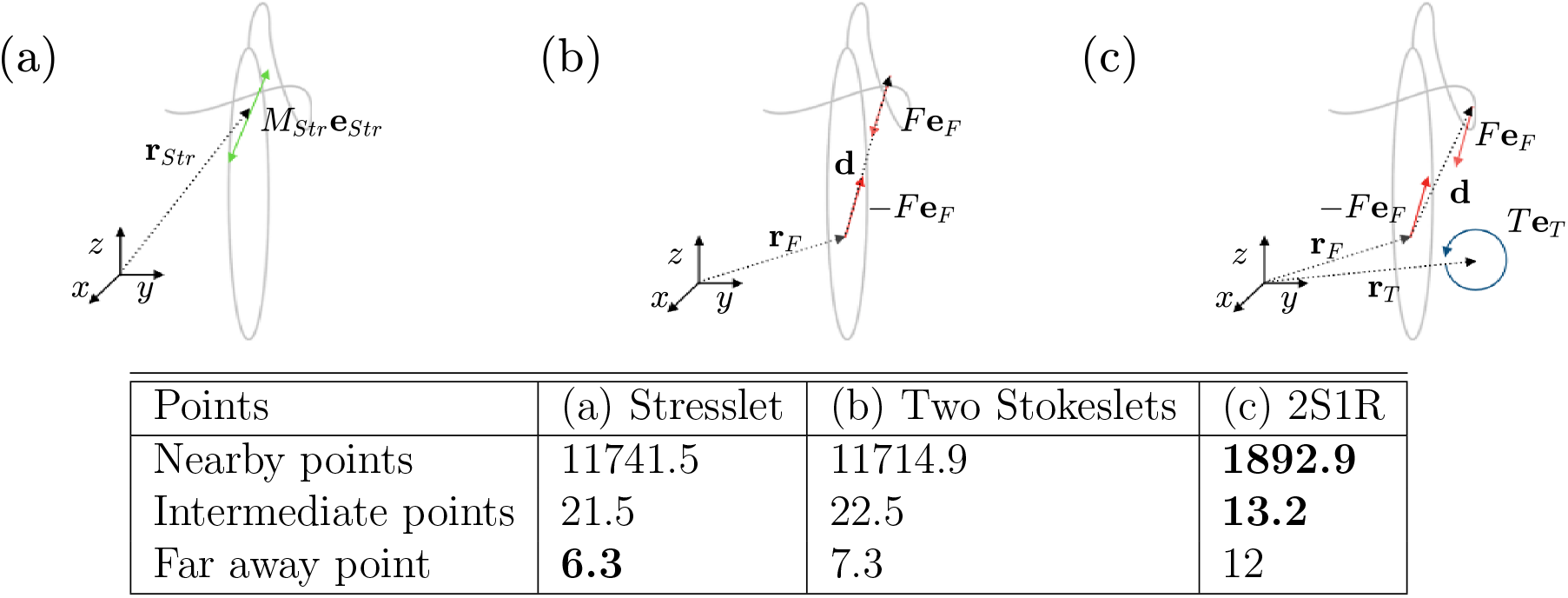
(a), (b) and (c) describe the three different singularity models we analyze for the reconstruction of the far field velocities: the Stresslet, the Stokeslet doublet, and the two Stokeslets one Rotlet model (2S1R). The table compares the Akaike Information Criterion for the three different approximations on nearby, intermediate, and far away points. We highlight in boldface the results corresponding to the best performing model.

As for the case of the *Euglena*, we first consider a Stresslet of magnitude *M*_*Str*_ directed along the unit vector **e**_*Str*_ and located at **r**_*Str*_. We define the set of parameter for the Stresslet as

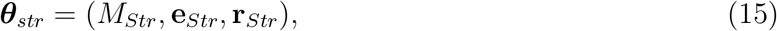

and choose values for these parameters in order to best match the flow velocities reconstructed in the previous sections, referred to as BEM reference velocities in what follows. We define **e**_*Str*_ as obtained from the unit vector **e**_*x*_ of the *x*-axis through the action of two rotation matrices, namely

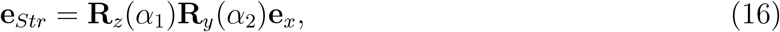

so that the two parameters *α*_1_ and *α*_2_ describe the orientation of the Stresslet. We rewrite the set of parameters for the Stresslet approximation as

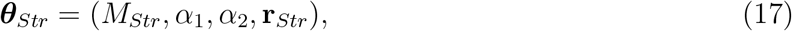

and write the fluid velocity at place **x** as

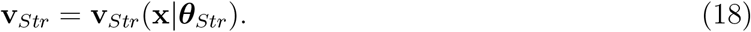

As for the approximation of the velocity field, we remark that this can be obtained either at any of the 10 instants *t*_*i*_ at which experimental results are available, or for the mean velocity during the stroke. The optimal set of parameters ***θ***_*Str*_ will vary accordingly. We find all the optimal approximations in the following, describing explicitly only the procedure for one of the 10 instants *t*_*i*_. To proceed, we define the error of the Stresslet approximation as the *L*_2_ norm of the difference between the reference BEM velocities and (18). Thus, we compute the optimal set of parameters through minimization of the error, that is

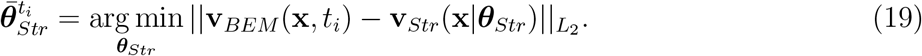

In particular, we use the global minimization algorithm introduced in [54] to solve the minimization problem of (19). The Stresslet is known to approximate the main features of the external flow on far away points but it fails to recover the flow field information near the cell body.

To achieve a better representation of the flow field we next introduce a different minimal system of singularities, inspired by the body kinematics of the swimmer, similarly to what has been done in [36] for *C. reinhardtii*. We use two opposite eccentric Stokeslet singularities, mimicking the propulsive force of the flagellum and the drag of the body, coupled with a Rotlet to balance the torque generated by the eccentricity of the forces. This system of two Stokeslets and one Rotlet (2S1R) includes as a special case the two Stokeslet approximation (if the forces are not eccentric) and also the Stresslet model analyzed above (in the limit of vanishing separation). These two new systems of singularities are shown in Table 1(b)-(c).

To parametrize the eccentric Stokeslet doublet, we use the direction of one Stokeslet **e**_*F*_, its magnitude *F* and position **r**_*F*_, and the relative position of the other Stokeslet **d**. As for the Rotlet, this is identified by its direction **e**_*T*_, magnitude *T* and position **r**_*T*_, so that the set of parameters for this approximation reads

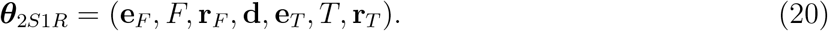

As previously done, we parametrize the orientation of the two Stokeslets as

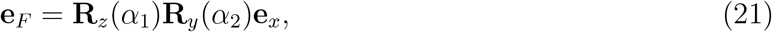

and impose the conservation of the angular momentum by requiring that

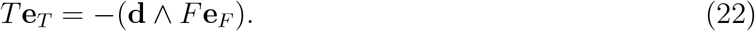

We rewrite the set of parameters for this approximation as

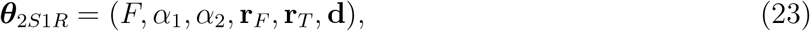

and define the induced velocity at place **x** as

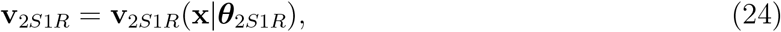

such that, proceeding as for (19), the optimal set of parameters is obtained by minimizing the error

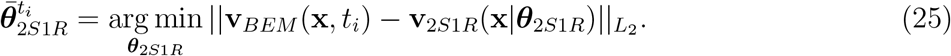

The results relative to the mean velocity during the stroke for the three different models on the three different spherical shells are shown in Table 1. We use the Akaike Information Criterion [55] to assess the quality of each model. We report in Table 1 the AIC with respect to the average velocity for the case of two eccentric Stokeslets and for points in the three spherical shells around the swimmer as defined above. We notice that the 2S1R model recovers a better statistical information on nearby and intermediate points. On far away points, the 2S1R model does not recover more information with respect to the Stresslet model. We also notice that the simple two Stokeslet model is equivalent to the stresslet as we can expect.

We now extend our analysis to the geometry of the flow field in close proximity to the cell body, in order to test whether the singularity models provide a realistic representation of the forces that *E. gracilis* exerts on the fluid. Of course, we only expect a qualitative agreement since, in the near field, the flow is strongly influenced by the presence of body and flagellum, with their actual shapes, which are not present in the flow field generated by the singularities. Furthermore, Stokes singularities lead to large velocity gradients, that are not physical.

For the sake of clarity, we report in Figure 6 velocity fields in the *xz*-plane. Specifically, a comparison in terms of the mean velocity field between the 2S1R approximation (left column) and the BEM (right column) is shown in the first row of the figure. We notice a qualitative agreement in some features, such as the occurrence of significant velocities at the left hand side of the flagellum (where one Stokeslet is located). Also, the flow exhibits an off-axis puller signature (i.e., the axes of the flow are not aligned with the longitudinal axis of the body). Nevertheless, the two mean flows differ as a consequence of the coarse-graining due to the singularity approximation. In fact, the action of the flagellum is described by a single point force, with a significant loss of information in the mean velocity field. The 2S1R approximation also introduces a visible peak in the velocity field at the center of the cell body. The rationale behind it is twofold: (i) the torque resulting from the tangential forces induced by the rotations of the body is approximated by a concentrated Rotlet and (ii) the forces exerted by the flagellum are more complex than what can be captured by a single Stokeslet, with part of them contributing to the Stokeslet near the body center. The resulting effect is a strong singularity in the velocity field near the body center, with velocities having much higher magnitudes than those from the BEM.

**Figure 6:**
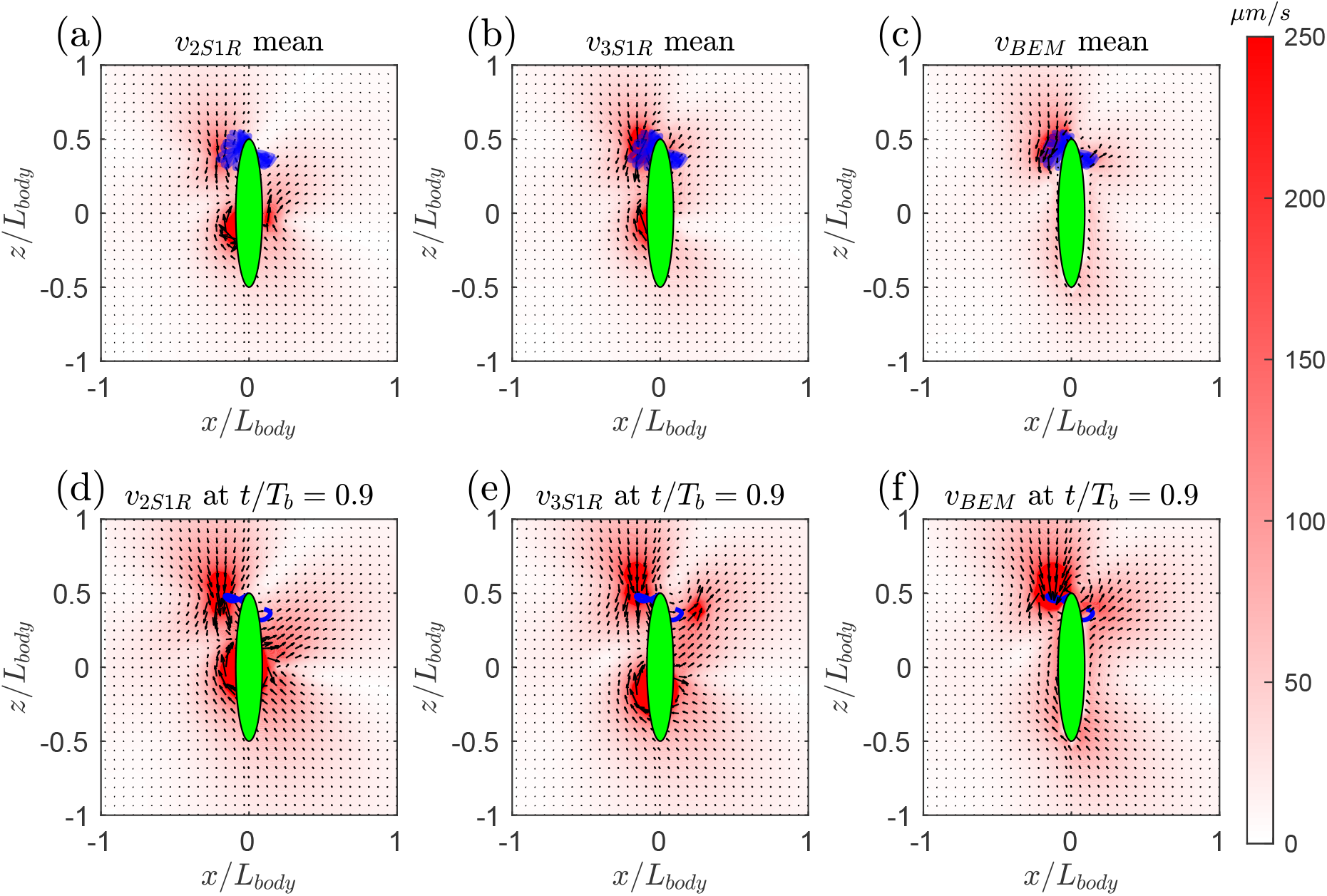
Qualitative flow field comparison in the *xz*-plane between the BEM (third column) and two singularity approximations: 2S1R used to approximate the far field (left column) and 3S1R (middle column). The first row represents the mean velocity, while the second reports the velocity computed for *t/T*_*b*_ = 0.9. The colormap represents the magnitude of the velocity field as projected on the *xz*-plane, while the quivers show the velocity projected on the plane. In all the figures, we superimpose in green the cell body and in blue the flagellar shapes. In particular, all the 10 flagellar shapes are shown in (a), (b) and (c), while only the flagellar shape relative to *t/T*_*b*_ = 0.9 is reported in (d), (e) and (f).

To better recover the flow field in the immediate proximity of the swimmer body, i.e. at a distance smaller than 1.5*L*_*body*_, we enrich the 2S1R approximation by using three Stokeslets and one Rotlet (3S1R). In view of the force and torque balance of (14), the three force singularities are such that

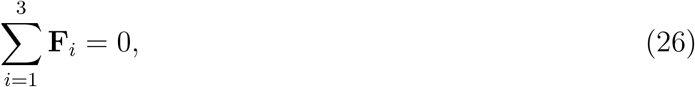

whereas the Rotlet has to guarantee the torque balance, that is

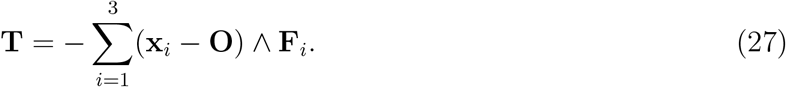

We optimize the corresponding velocity field **V**_3*S*1*R*_ = *V*_3*S*1*R*_(**x**|***θ***_3*S*1*R*_) to match results from the BEM on points between 0.75*L*_*body*_ and 1.5*L*_*body*_, and obtain a reduction of the error of more than 50% with respect to the 2S1R approximation. Results for the 3S1R model are shown at the center of Figure 6 and highlight that two point forces are exploited by this approximation to model the action of the flagellum on the fluid.

This fact is further emphasized by Figure 7, where the 2S1R (left) and the 3S1R (right) models are compared. In particular, the figure shows the respective systems of forces as superimposed to the traction field **f** from the BEM. Interestingly, we notice that an increase in the complexity of the model in terms of number of singularities leads to a more accurate representation of the continuous traction field. In passing, we notice that a better approximation in the near field could be achieved by further increasing the number of singularities, although at the expense of increasing the number of parameters to account for in modelling the system.

**Figure 7:**
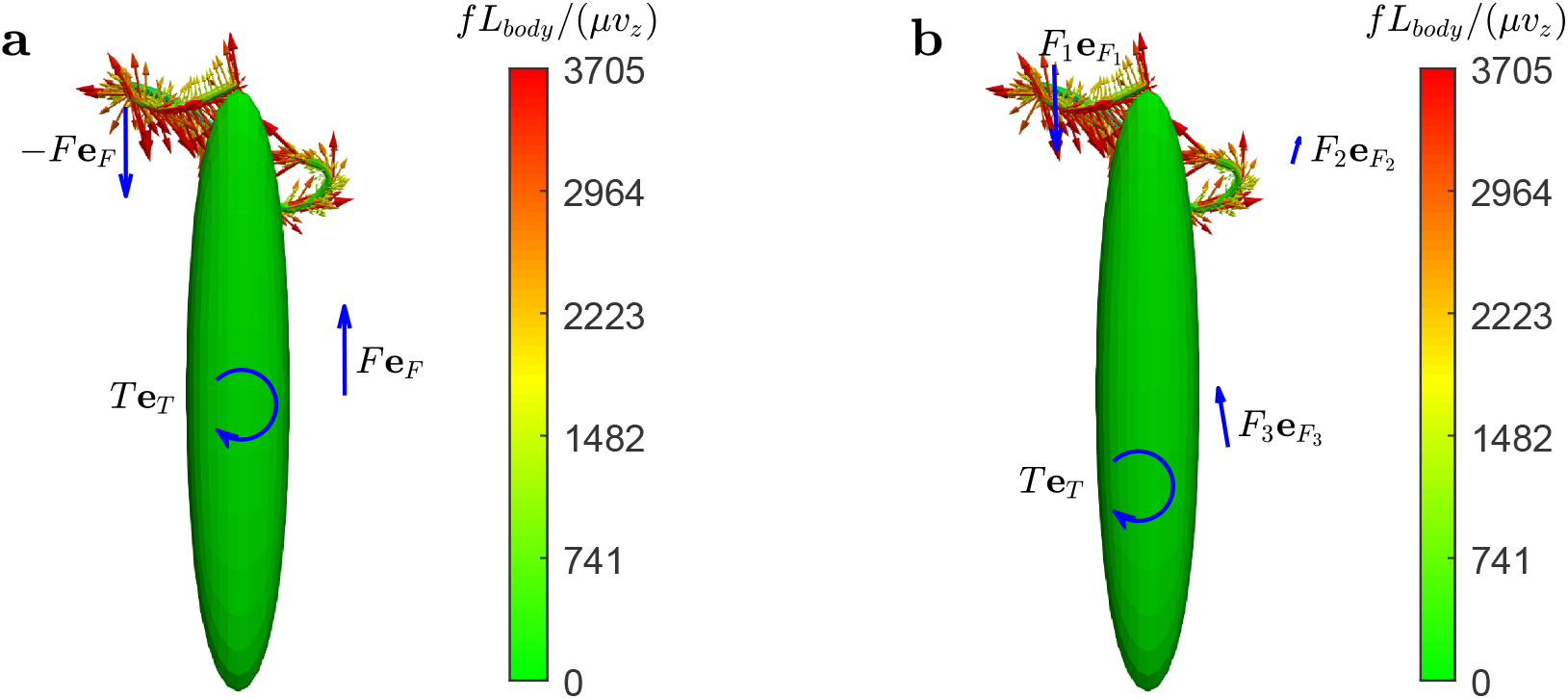
Qualitative comparison in the *xz*-plane between the BEM tractions and the singularity systems 2S1R (a) and 3S1R (b). The torques represented in blue are projections along the *y* axis. Blue arrows on the *xz*-plane complement the representation of the singularity systems. The colormap is for the magnitude *f* = |**f**| of the BEM tractions **f** on the *xz*-plane.

To prove the effectiveness of the qualitative reconstruction in the near field, we analyze in the second row of Figure 6 the flow field at time *t/T*_*b*_ = 0.9. This exhibits a clear puller signature with, in addition, rotational flows induced by the eccentricity of the flagellum. Both these facts are recovered by the two singularity approximations. Only the 3S1R model, however, is able to capture a feature revealed by the BEM analysis, namely, that the forces exerted by the flagellum are large and propulsive (directed downwards) near the proximal end, while they are smaller but resistive (directed upwards) at the distal end. This is clearly visible in Figure 7, where we recall that the tractions exerted by the flagellum on the fluid, as computed using the BEM, are compared with the two systems of singularities described above. Returning to Figure 6, we still recognize a significant peak in the velocity magnitude at the center of the body. As discussed earlier, this is induced by having concentrated at a singular point all the distributed forces exerted by the cell body.

From our analysis, we conclude that the 2S1R approximation provides a satisfactory way to rationalize the flow field induced by a sample of swimming *E. gracilis* at distances larger than 1.5*L*_*body*_. To arrive at a better approximation in the immediate proximity of the cell body we had to account for more singularities to capture the complexity of the action of the flagellum on the surrounding fluid, as demonstrated by the 3S1R approximation.

### 4.2 Characterization of the swimming behavior using multipole expansion

A different approach to study the swimming behavior of micro-swimmers is to consider the leading order term of the multipole expansion corresponding to the boundary integral equation. Following [40] and [18], we introduce the dipole matrix **D**, which represents the leading order term of the multipole expansion for the operators appearing in the representation formula of the Stokes system,

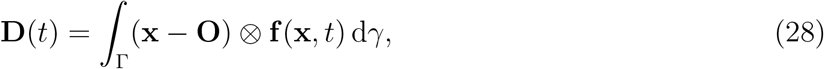

where **f** is the traction acting at place **x** of the boundary Γ of the swimmer’s body. We then introduce the Stresslet matrix **S** as

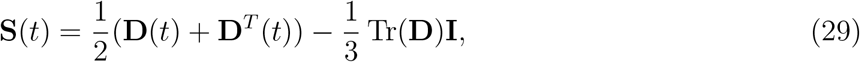

which can be computed by means of the BEM. As explained in [18], the determination of whether a swimmer is of pusher or puller type follows from the sign of the scalar quantity

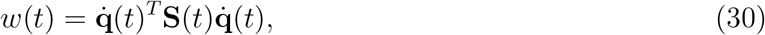

in which 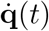 is the velocity of the swimmer at time *t*. More explicitly, a negative sign of *w* identifies a puller, whereas a positive sign of *w* corresponds to a pusher.

By applying the criterion above with reference to tractions and velocities computed with the BEM, we find that *E. gracilis* is a micro-swimmer of mixed type during a stroke, a fact in agreement with the observation of mixed pusher-puller behaviors in complex eukaryotic swimmers, as reported in [18]. Specifically, we compute a mean value for *w*(*t*) of 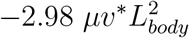, while the maximum and minimum values are 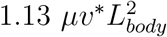 and 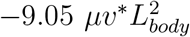, respectively. Here *v*^*^ denotes the absolute value of the mean velocity, so that, on average, *E. gracilis* behaves as a swimmer of puller type. We then consider the mean of the Stresslet matrix **S**(*t*) during a stroke and compute its eigenvector corresponding to the eigenvalue of largest absolute value. We notice that this is not aligned with the longitudinal axis of the swimmer, thus confirming that *E. gracilis* is an off-axis puller for what concerns the mean flow.

## 5 Conclusions

The analysis of the flow fields induced in the surrounding fluid offers insight on the swimming behavior of micro-organisms. In this work we have presented the first numerical reconstruction of the three-dimensional flow field generated by a specimen of *E. gracilis* swimming using the “spinning lasso” flagellar mechanism, a non-planar waveform. We have accomplished this by reconstructing the history of flagellar shapes and shape velocities, and then computing the resulting flows by solving numerically the Stokes equations with a Boundary Element Method. The experimentally measured translational and angular velocities of the swimmer, and its observed trajectories are reproduced very accurately by our procedure. This fact provides a strong argument supporting our reconstruction of the swimming stroke.

Furthermore, we have validated our numerical reconstruction of the flow field against experimental measurements obtained with the General Defocusing Particle Tracking method. This technique allows to track the three-dimensional position of passive tracer particles dispersed in the fluid, and hence to reconstruct the complete flow field. This is needed here since *E. gracilis* uses a non-planar flagellar beat. Looking at the time- and azimuthally-averaged velocity fields, we could identify three characteristic flow structures for all the three velocity components, both in the numerical and in the experimental data. The good agreement between experimental data and numerical results justifies the use of the numerical model (our synthetic stroke) to analyze in more detail the swimmer mechanism of real *Euglena* cells.

We have then analyzed the flow fields (computed by solving numerically the Stokes equations via BEM, using as input our synthetic stroke), and coarse-grained them in terms of a few dominant Stokes flow singularities. We find that a system of two eccentric Stokeslets and one Rotlet is able to approximate well the velocity field away from the body, and captures some of the main features of the swimming mechanism of the organism. Moreover, the BEM analysis shows that the flagellum may exert forces that change orientation along its length (e.g., propulsive forces near the proximal end and resistive ones at the distal end). To capture this effect and, more generally, to represent adequately the signature on the generated flow left by flagella and cilia that do not beat in a single plane, more complex systems of singularities would be needed. We hope that our study may motivate also others to investigate the question of which systems of singularities may be effective in coarse-graining the flows associated with the three-dimensional trajectories that have attracted interest in the recent bio-physical literature on micro-swimmers [8, 13, 24, 25, 26, 27, 28].

Finally, by combining the singularity analysis with a multipole expansion we conclude that, on average during one stroke, *E. gracilis* can be idealized as an off-axis puller. Its instantaneous character during the stroke oscillates between a puller and a pusher behavior, similarly to what has been observed for other micro-organisms such as sperm cells and *C. reinhardtii*.

## Acknowledgements

This work has been supported by the ERC Advanced Grant 340685 MicroMotility. M.R. acknowledges funding from the European Union’s Horizon 2020 research and innovation programme under the Marie Sklodowska-Curie grant agreement no. 713683 (COFUNDfellowsDTU).

