## Supplemental material for "How *Euglena gracilis* swims: flow field reconstruction and analysis"

### Supplementary Material for How *Euglena gracilis* swims: flow field reconstruction and analysis

#### I Boundary Element Method

In this section, we briefly describe the numerical procedure to find the rigid velocities $\dot{\mathbf{q}}$ and $\boldsymbol{\omega}$ appearing in (2), and characterizing the translational and rotational motion of the swimmer, once the current configuration of the swimmer and its shape velocities $\dot{s}$ are assigned.

We discretize the shape velocities $\dot{s}$ and the tractions $\mathbf{f}$, and describe the latter using $N$ scalar unknowns. We write the equations relating the velocity $\dot{\mathbf{x}}$ (which, from (2), depend on the unknown rigid velocities $\dot{\mathbf{q}}$ and $\boldsymbol{\omega}$, and on the given discretized shape velocities) to tractions $\mathbf{f}$ at $N$ distinct points (i.e., we use a collocation method to solve the Boundary Integral Equation (BIE) associated with the Stokes equations (6), see [1, 2]). In addition, we write the six balance laws of linear and angular momentum, equations (3). We define in this way a system of $N+6$ equations in $N+6$ unknowns which we solve numerically to obtain the discrete representation of the traction forces ($N$ degrees of freedom) and the 6 unknown components of the rigid velocities $\dot{\mathbf{p}}(t) = (\dot{\mathbf{q}}(t), \boldsymbol{\omega}(t))$.

A few additional details on our solution strategies are given in what follows. The numerical procedure to solve a BIE leads to a Boundary Element Method (BEM). Among the several different implementations available in the literature (see, e.g., [3] and the references cited therein) we follow the solution scheme described in [4, 5]. The resulting BEM [6] exploits distributed memory parallelism (MPI) and couples together existing OpenSOURCE High Performance Computing libraries. The final discretization consists of 384 cells for the cell body and 648 for the flagellum. Figure 2 i the main text shows the numerical domain obtained using standard Lagrangian linear element (the corresponding discrete representation consists of 1032 cells and consequently $N = 3504$ unknowns for the vector unknown for $\mathbf{f}$).

We use a classic collocation scheme to derive the final linear system: we replace the continuous functions for the velocity $\dot{\mathbf{x}}$ and for the tractions $\mathbf{f}$ with their finite element approximations and we satisfy the corresponding discretized equation on a number of points equal to the number of unknowns.

Moving now to the concrete implementation of this algorithm to the case of a swimming cell, we use the kinematics described in [7] for a swimming specimen of *E. gracilis*. We approximate the cell body with a rigid prolate spheroid having major axis of length 25.14 µm and minor axes of length

4.345 μm, and we describe the flagellum as a morphing elongated spheroid having a cross sectional diameter of 0.75 μm. In particular, we superimpose the spheroid centerline to the experimental observations of [7]. We have in this way experimental data for 10 different flagellar shapes of an *E. gracilis* cell during one stroke. In order to determine its rigid velocities $\dot{\mathbf{q}}$ and $\boldsymbol{\omega}$ using the algorithm described in this section, we would need to know the shape velocities $\dot{s}$ of the flagellum at each of the 10 different shapes. The experimental observations of [7] do not provide this information. Section 2.2 shows how we have overcome this problem.

It is possible to use the same BEM procedure to compute the velocity field $\mathbf{v}$ at any place of the fluid domain $\mathbb{R}^3 \setminus B_t$. An open source numerical code that performs this calculation can be found in [8].

#### II Experimental setup

All the experimental observations were performed on specimens of *E. gracilis* (strain SAG 1224-5-27) obtained from the SAG Culture Collection of Algae at the University of Göttingen. Samples were maintained axenic in liquid culture medium Eg and cultures were transferred weekly. For each experimental trial, a dilute solution of *E. gracilis* and fluorescent polystyrene beads with diameter of 1 μm (Life Technologies, catalog number F8821) in culture medium was prepared and introduced between two microscope slides separated by a ∼ 80 μm thick double-sided adhesive spacer, following the procedure described in [7]. The fluorescent beads, with a final volume fraction of 0.06 %, were used as passive tracers for the PTV measurements. Swimming *Euglena* cells and beads were imaged in bright-field by using an Olympus IX 81 inverted microscope equipped with a LCAch 40 X Ph2 objective (numerical aperture 0.55) and exploiting the built-in magnification changer of 1.6 X. To implement the GDPT method and enhance the defocusing, a cylindrical lens with focal length of 1000 mm was fixed along the imaging path in front of the microscope camera port to introduce a mild astigmatic aberration [9, 10]. Several movies of swimming *E. gracilis* were recorded at a frame rate of 1000 fps using a high-speed complementary metal–oxide semiconductor digital camera (Photron, model FASTCAM Mini UX100). The focus of the objective was adjusted in order to observe only specimens swimming parallel to the mid-plane of the chamber. In this way, the bottom and top walls are far enough from the cell body and have a negligible effect on the flow field, as highlighted in Section 2.2 of the manuscript. Among the image recordings, only those in which the specimen was swimming regularly were selected. A specimen was considered to swim regularly, if it maintained a steady flagellar stroke across the recording (tipically 20-30 beats), resulting in a helical trajectory along a straight screw axis. A thorough analysis of the variance of the flagellar beating of the same and different specimens on regular swimming was performed in [7]. Such analysis showed that, although some variability in the kinematic parameters emerged, all specimens shared the same qualitative swimming geometries. The procedure described in [7] was used to reconstruct the helical trajectories of the swimming specimens and the relative orientation in space.

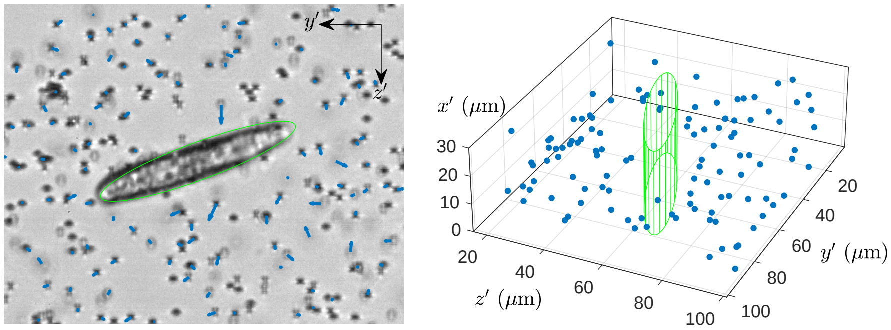


**Movie S1** Experimental images of the swimming *Euglena* with the tracer particles (1-μm-diameter polystyrene spheres) and corresponding three-dimensional particle positions obtained from GDPT measurements.

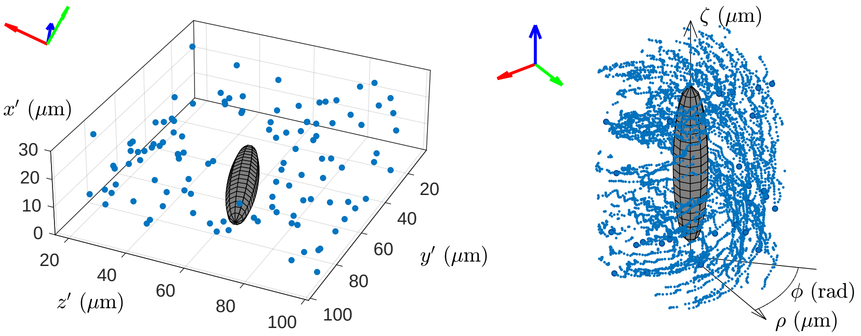


**Movie S2** Reconstructed position of the swimmer and tracer particles as observed in the laboratory reference frame (left) and in the body reference frame in cylndrical coordinates (right).
